# Ambient humidity and temperature influence physicochemical drift during laboratory storage of field-collected mosquito breeding water

**DOI:** 10.64898/2026.04.10.717870

**Authors:** Jeffrey K. Boateng, Bernard Agyei Adams, Fred Aboagye-Antwi, Jewelna Akorli

## Abstract

The use of field water for laboratory rearing of mosquitoes could offer a better representation of the natural aquatic environment than laboratory tap or deionised water. For logistical reasons, such water may be stored in the laboratory environment for an extended period, but its stability is poorly documented. This study evaluated the influence of laboratory storage conditions on the kinetics of physicochemical parameters of breeding water collected from a field habitat. To capture within-habitat variability, water was collected from multiple spatial points from a breeding site and transferred into plastic containers for storage under laboratory conditions. Water physicochemical parameters were measured in the field to establish baseline readings, while laboratory measurements were done at 2-3-day intervals over 2 months to evaluate temporal changes. A linear mixed-effects model was fitted to evaluate the determinants of changes in physicochemical parameters under laboratory storage. Most parameters exhibited high stability; however, water temperature increased significantly by an average of ∼1.5℃ (*p*= 0.046) relative to the field. Water pH demonstrated a long-term rise over the 2-month storage period with a transient, significant dip of 0.71 units after a week of storage (*p*< 0.001). Overall, LMM analyses revealed that ambient relative humidity was the strongest statistical predictor of change in all water parameters except pH (*p*< 0.05). Ambient temperature correlated positively with water temperature and ammonium nitrogen (NH_4_-N) (*p*<0.002), and negatively with dissolved oxygen (*p*< 0.002). These results indicate that stored field water is highly sensitive to the laboratory microclimate. Specifically, water temperature, pH, and NH_4_-N serve as candidate indicators for storage-related physicochemical drift. We recommend the rigorous standardisation of insectary humidity and temperature, and monitoring of water parameters, which are likely relevant for bioassay reproducibility.

## INTRODUCTION

The maintenance of mosquito populations in the insectary has been a common practice for vector research since the beginning of the 20^th^ century, when *Aedes aegypti* was first colonised in the laboratory. Eggs are hatched either in deionised water mixed with nutrient broth and yeast or deoxygenated boiled distilled water with the addition of larval diet^1^. In standard insectary practice for mosquito management, dechlorinated water is commonly used for hatching eggs and maintaining larvae^2^. However, depending on the objectives of one’s study, water could be sourced from breeding ponds in the field for maintenance of mosquitoes in the insectary; for example, to conserve field-acquired midgut microbiota in mosquitoes during laboratory colonization^3^. While laboratory experiments may employ the use of water collected from the field *ad libitum*, this is logistically challenging and may necessitate storing these field-sourced water samples for periods ranging from hours to several weeks before use. The validity of this resource depends largely on maintaining the *in-situ* characteristics of the collected water to ensure experimental stability. Water parameters including pH, conductivity, turbidity, dissolved oxygen, temperature, nutrient concentrations (nitrogen and phosphorus) and microbial community composition, can influence larval ecology^4^, vector competence, fitness and adult population dynamics^5,6^.

Numerous studies have been conducted on the water quality assessment of mosquito breeding habitats ^7–11^, however, far less attention has been given to assessing the temporal stability of water under laboratory storage conditions. Storage may initiate a cascade of chemical and biological shifts. For example, organic compounds that serve as oviposition cues can evaporate within hours of collection, directly affecting behavioural assays and population-level responses^11–13^. More broadly, microbial communities which serve as food sources for larvae, undergo compositional drifts during storage, altering nutrient cycling and availability, dissolved gas production and mineral concentrations^14,15^. While studies have shown substantial shifts in microbial communities and dissolved components in stored samples over short timescales^16,17^, the kinetics and drivers of these drifts remain a critical blind spot in laboratory practice.

This study employed an observational, longitudinal design to monitor temporal changes in selected physicochemical parameters in field-sourced mosquito breeding water stored under standard laboratory conditions to provide evidence-based recommendations for breeding water storage in mosquito research. We monitored parameters critical to mosquito development in laboratory breeding including temperature, which is the main driver of metabolic rate and development; pH which affects enzyme function and ion balance; dissolved oxygen which is essential for aerobic respiration and sediment redox status; conductivity which is an indicator of ionic strength and influences osmoregulation; and total dissolved solids which is a measure of the organic matter available in the water^18–20^. The selected parameters represent key physicochemical factors influencing mosquito larval habitat suitability, and were prioritized to assess their stability during laboratory storage rather than to provide a full ecological characterization of the breeding water.

## MATERIALS AND METHODS

### In situ measurement of water parameters and water collection

Water samples were collected from a mosquito breeding pond on an urban agricultural site popular for several reported mosquito-related field activities conducted in Accra, Ghana^21–27^. At the time of collection, rainfall was scanty and dugouts that usually held water were either dry, held too little water, or had no mosquito larvae or pupae. One breeding site with dimensions of about 5m × 3m (5°35′55.5″ N, 0°10′52.9″ W) was identified based on the evidence of *Anopheles* mosquito larval activity.

To capture spatial heterogeneity, three sampling points (A, B, C) were established in a triangular configuration across the circumference of the selected breeding pond. Baseline estimates of physicochemical parameters were measured *in situ* using a YSI ProQuatro Multiparameter meter (YSI Inc., USA). Before baseline measurements, the multiparameter probe was calibrated using standard solutions according to the manufacturer’s protocol. Parameters of interest included temperature, pH, conductivity (COND), total dissolved solids (TDS), salinity (SAL), dissolved oxygen (DO), and ammonium-nitrogen (NH_4_-N). All readings were taken in triplicate at a depth of 10-15cm, and the multiparameter probe was rinsed with dechlorinated tap water and blotted dry between each replicate to prevent cross-contamination.

Following *in situ* measurements, three 25L high-density polyethene (HDPE) plastic containers (two green and one yellow) were rinsed with water from their respective sampling points to wash out any previous residue and prevent cross-contamination, then filled to about 90% with their respective water samples. The containers were sealed with airtight lids and transferred to the laboratory.

### Monitoring of water samples in the insectary

In the laboratory, the storage containers were kept closed and placed side-by-side in a climate-controlled insectary maintained at 27 ± 2°C temperature, 75 ± 10% relative humidity (RH) and 12:12h light: dark photoperiod. The physicochemical parameters were measured twice weekly for two months to assess temporal dynamics under storage conditions. Before each measurement, storage containers were gently swirled to resuspend any settled material and 500ml of the stored water was then poured into open larval rearing trays (L35× W26× H4.5 cm) to mimic air-water surface exposure typical of standard mosquito rearing conditions. Physicochemical measurements were taken in triplicate immediately using a multiparameter probe, after which the water was returned to its respective container, thus, the study evaluated the stability of breeding water parameters during laboratory storage under routine monitoring conditions rather than under completely undisturbed storage Triplicate probe readings at each time point within each container were technical replicates and were averaged before inferential analysis. Insectary conditions were also monitored alongside using the HOBO^®^ temp/RH logger (Onset Corporation SE, USA) during the study period, and the data were retrieved via the HOBO^®^ Connect application software.

### Data analyses

All triplicate measurements were averaged to obtain a single representative value per parameter for each container and time point before all statistical analyses. Descriptive analysis was used to summarise physicochemical estimates and variability in breeding water collected in the field and stored in the laboratory for each parameter and continuous variables were summarised as median and interquartile range (IQR). Linear mixed-effect models (LMMs) were used as the primary analytical tool to account for the longitudinal, nested structure of the data (repeated measures within-pond replicates) while evaluating the determinants of changes in physicochemical parameters. Days since collection (time), ambient insectary temperature and relative humidity were treated as fixed explanatory variables, while container (sample ID) was included as a random effect. To identify the specific days on which laboratory water parameters diverged from initial field conditions, a “Many-to-one” framework embedded within LMM was employed. For each parameter, “Day” was treated as a categorical fixed effect with initial collection (Day 0) set as the reference level to allow for direct comparison between subsequent storage days and baseline readings. To account for temporal autocorrelation arising from repeated measurements from the same containers over time, a first-order autoregressive [AR (1)] correlation structure was included. Model outputs were evaluated through visual inspection of residual plots to assess the direction and magnitude of effects, and identify parameters that exhibited coordinated changes during storage.

Locally estimated scatterplot smoothing (LOESS) curves were used to illustrate overall temporal trends and gradual shifts in physicochemical parameters of the breeding water during laboratory storage. Spearman’s rank correlation plot was used to visualise the strength and direction of associations among physicochemical parameters, providing context for the linear modelling results and highlighting coordinated changes that may not be captured by individual model coefficients. All statistical analyses were performed in R version 4.5.2 via R Studio (2025.05.0+496) using custom scripts.

## RESULTS

### Temporal changes in water parameters under laboratory conditions

Over the 2-month monitoring period, we took readings of the water in each stored container in triplicate. These were averaged as the overall laboratory state of the water and compared to the baseline field water estimates. Overall, temperature and dissolved oxygen varied significantly between field and lab-stored water, with temperature increasing by an average of ∼1.5℃ (*p*< 0.001) over the storage time (Fig. 1). Although steady and continuous change for most physicochemical parameters was not observed in the stored water, there were notable time point fluctuations that differed from the field water conditions (Fig 2, Table S1). Dissolved oxygen exhibited an immediate decline by Day 3 (-21.6mg/L, *p*= 0.0005) of storage and remained stable at that concentration (*p*= 0.27) thereafter. There was significant accumulation of ammonium nitrogen (NH_4_-N) on Day 15 and Day 24 (0.08 – 0.12mg/L, *p*< 0.05), though levels recovered to field water estimates. The water showed an overall cumulative increase in temperature of 0.87℃ from Day 3 to 1.71℃ by Day 45 of storage (*p*< 0.001) (Table S1). Similarly, pH showed a long-term rise (*p*= 0.02) with a transient, significant dip of -0.71 on Day 7 (*p*< 0.001).

**Fig. 1:**
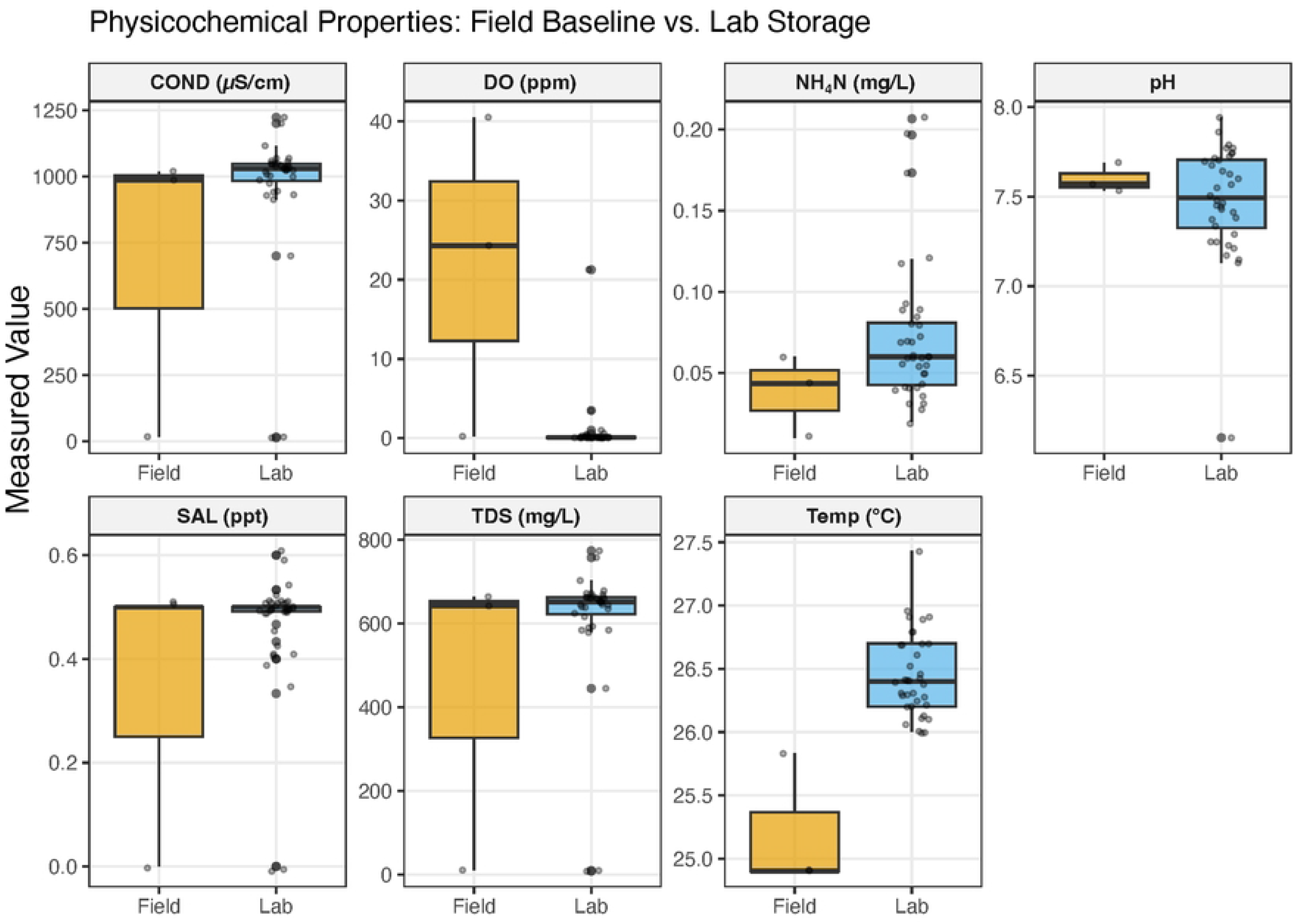
Comparison of physicochemical parameters between field-collected and laboratory-stored mosquito breeding water. Grey dots represent the aggregated daily measurements from sources over the monitoring period. Box size represents the interquartile range of the distribution of values of each parameter, and horizontal lines within the boxes are medians.

**Fig. 2:**
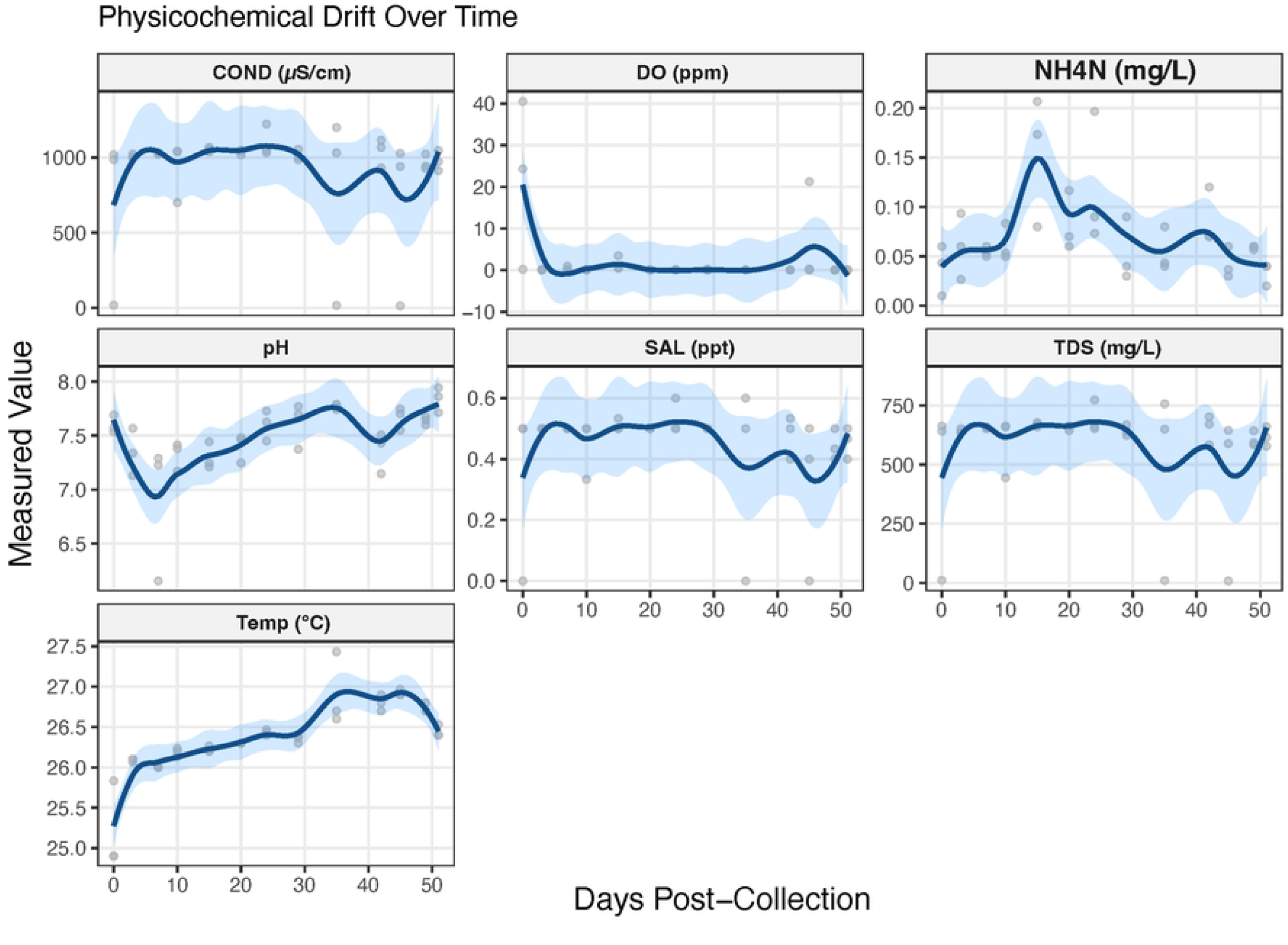
Temporal trends in water parameters during laboratory storage. Blue line represents LOESS-smoothed trends with 95% confidence intervals (light blue shading). Points indicate average measurements of parameters in each pond water during monitoring period.

### Relationship between insectary environmental conditions and water parameters

We also recorded insectary temperature and relative humidity over the monitoring period. Laboratory environmental conditions fluctuated over the study period, with temperature and relative humidity reaching an average of 26.8 ℃ and 83.1%, respectively, and varying inversely over time (Fig. S1). A linear mixed-effects model (LMM) was fitted to combine insectary and water data to explore the influence of the environmental parameters and days on stored water physicochemical conditions (Fig 3, Table S2). Ambient relative humidity was a significant driver of change in all water parameters except pH (Fig 3A, Table S2). It correlated positively with conductivity (COND), ammonium nitrogen (NH_4_-N), salinity (SAL) and total dissolved solids (TDS) and, negatively with water temperature and dissolved oxygen (DO) (Fig 3B). As expected, environmental temperature correlated positively (r= 0.75, *p*<0.0001) with water temperature indicating water equilibration to the insectary temperature. A 1℃ increase in ambient temperature also influenced a 0.12mg/L rise in NH_4_-N (*p*= 0.001), and 15.8ppm (*p*= 0.02) decrease in DO. Storage days most influenced water temperature, pH and NH_4_N (Fig 3, Table S2); NH_4_N decreased subtly with prolonged water storage.

**Fig 3:**
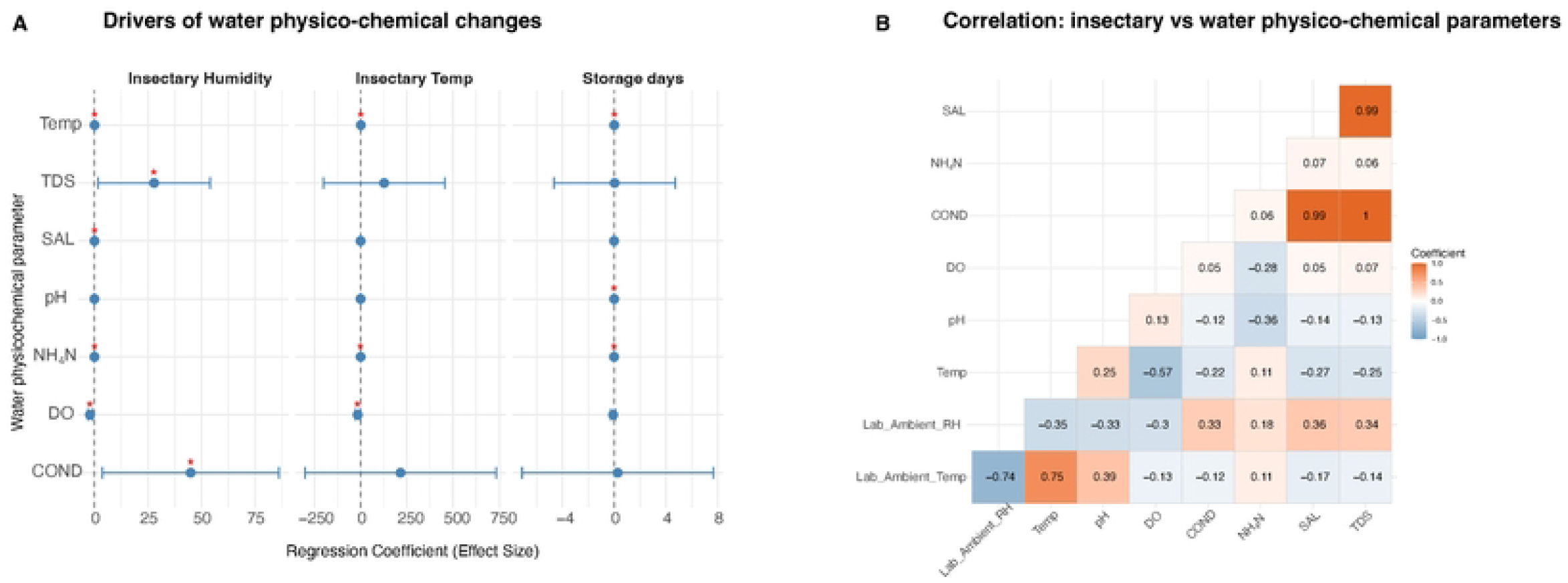
Determinants of changes in water physico-chemical parameters and correlations with insectary environmental conditions. The laboratory ambient temperature (℃), relative humidity (%) and length of storage days were set as independent factors for the linear mixed-effects model (A). Points show estimates (regression coefficient) with blue horizontal bars as 95% CI; red asterisk (*) indicates *p* < 0.05. Correlation plot shows strength of the relationship between and among insectary conditions and water parameters (B).

## DISCUSSION

The role of the aquatic environment in shaping the physiology of mosquito vectors is important for understanding vector biology and identifying environmentally sustainable vector control strategies. Laboratory studies that leverage field water or cryopreservation of field-derived microcosms could allow more relevant laboratory-to-field interpretation^28–30^. We have studied the changes in field water parameters during their storage in laboratory conditions. Our results showed that the most dramatic change occurred in dissolved oxygen (DO), which could be due to movement of water from the field to laboratory conditions rather than to how long the water was stored or to measurement perturbation. This reduction could also be associated with limited atmospheric gas exchange in the enclosed plastic container during laboratory storage, unlike water in the field, which remains exposed to the environment, allowing for higher oxygen availability and more active microbial turnover. Mosquito larvae primarily obtain oxygen from the atmosphere through their spiracles, but can also utilise oxygen directly from water^31,32^. Consequently, lower DO levels in stored water may impose additional physiological stress on developing larvae. This highlights the importance of maintaining adequate aeration during laboratory storage to better mimic field conditions and reduce potential developmental stress.

On the contrary, water temperature, pH and ammonium nitrogen levels were significantly influenced by storage days. In addition, the ambient temperature and relative humidity were also significant determinants of water temperature and ammonium nitrogen, but not pH. Ambient conditions positively correlated with the thermal properties of the stored water^33,34^. The observed increase in pH and decline in ammonium concentrations under anaerobic storage likely reflect microbially mediated nitrogen transformation and proton-consuming reduction processes, resulting in gradual alkalinization of the stored water. Ammonium nitrogen (NH_4_-N) is a source of nutrients and substrate for nitrifying bacteria, which are abundant in environmental water and soil. Thus, a decline in NH_4_-N in aged water in response to laboratory temperature and humidity may reflect loss of metabolically active microorganisms^35^. This is crucial for field-derived mosquitoes if the research interests are in maintaining the vector’s natural gut microbiota. The microbial communities in the mosquito breeding water are ecologically important food sources for larvae and modify water chemistry through metabolic activities^36,37^. While these parameters are known to significantly affect *Anopheles* larval development and microbial diversity^38–41^, the values we recorded during the water storage still fall within acceptable ranges for successful mosquito larval breeding ^42,43^. Since microbial composition and larval performance were not measured, these biological interpretations remain provisional.

While our results provide good baseline information on the effects of laboratory microclimates on stored water, this study was confined to a single pond habitat, sampled at three within-site points with repeated measurements over time, which may limit the broader spatial representativeness of the findings. Another key limitation was that aliquots from the baseline collection and subsequent monitoring were not preserved for parallel microbial assessments or larval performance assays. If the stored water had been used for larval maintenance in the insectary, the addition of larval food, larval metabolism, and food decomposition may alter the parameters from those observed in the “resting” water, requiring direct evaluation of the performance of the stored water when in use. However, establishing this baseline for the stored water is a necessary precursor to understanding more complex, bio-loaded rearing systems.

Future studies should evaluate whether water with different organic loads or chemical compositions exhibits similar buffering capacities over time for experimental microclimate variations. While we used the same type and size of containers, the shape of the container used could have affected the rate of evaporation and gas exchange. Different storage vessels could be compared with open versus closed-container setups to investigate most appropriate holding conditions.

## CONCLUSION

This study demonstrates that the physicochemical stability of field-collected water is a result of both intrinsic temporal drift and extrinsic environmental modulation. The steady increase in alkalinity (pH) is the most consistent marker of water “age”. Physicochemical parameters were highly sensitive to daily fluctuations in insectary humidity, suggesting that changes in the mineral concentrations often attributed to water ageing, are actually a reflection of the facility’s climate control precision. Our data suggest that a candidate equilibration period may reduce early instability in DO, but this recommendation requires validation in larval performance or bioassay studies while maintaining stable insectary microclimates, particularly relative humidity, which showed strong statistical association with several water parameters. Routine monitoring and standardisation of storage conditions when using laboratory-stored field water in mosquito research are recommended for experimental reproducibility and consistency.

## SUPPORTING FILES

**Fig. S1: Environmental readings of the insectary during the study period**. Each dot represents the daily mean of measurements recorded at 15-minute intervals by the data logger.

**Fig S2. Trends in environmental conditions and physicochemical parameters during laboratory storage**. Panels show the trends between environmental temperature and (A) water temperature, (B) ammonium nitrogen and (F) dissolved oxygen. Panels (C, D, E) show the trends between relative humidity and water temperature (C), ammonium nitrogen (D) and dissolved oxygen (E). Dots represent aggregated values while lines indicate fitted trend curves.

**Table S1: Significant results of Dunn’s test for daily comparison of parameters in field vs lab stored water**.

**Table S2: Results of linear mixed effect model testing the influence of insectary environmental conditions on water parameters over the storage period**. Environmental drivers (ambient tempaerature and relative humidity) were set as fixed effects, with sample ID fitted as a random intercept (1|sample ID) to account for repeated measurements. Significant p-values are boldened.

## ACKNOWLEDGEMENTS

We thank the farmers of the agricultural site in Opeibea, Accra where the water samples were collected for their continuous support in making their farms available to us in the conduct of our vector research.

## AUTHOR CONTRIBUTIONS

JA and JKB conceptualised and designed the study. JKB and BAA conducted the research. JA and FA-A provided resources and instrumentation for the study. JA acquired funding and managed the project. JA and JKB performed formal analyses and visualisation of results. JKB wrote the original draft. JA reviewed and edited the draft for publication.

